# Cooperative antibiotic response in coupled biofilm and planktonic *E. faecalis* communities

**DOI:** 10.64898/2026.05.18.725849

**Authors:** Gabriela Fernandes Martins, Keanu Alexander Guardiola-Flores, Luis Zaman, Jordan M. Horowitz, Kelsey M. Hallinen, Kevin Wood

## Abstract

Bacterial communities grow as dynamic populations that respond to their environments. A clinically relevant example is the inactivation of beta-lactam antibiotics by intracellular beta-lactamase in *E. faecalis* resistant strains. In these populations, resistant bacteria act as antibiotic sinks, detoxifying the environment and allowing sensitive bacteria to survive treatment through a cooperative interaction. In this work, we study strongly coupled planktonic and biofilm populations of mixed sensitive-resistant *E. faecalis* bacteria under antibiotic stress using fluorescent microscopy. The presence of resistant bacteria in the system benefits both resistant and sensitive cells, leading to mixed planktonic and biofilm populations at super-inhibitory drug concentrations. We show that a beta-lactam antibiotic with or without the addition of a beta-lactam inhibitor can lead to a population inversion effect, characterized by a non-monotonic relation between initial and final fractions of resistant bacteria. The effect is observed in both the planktonic and biofilm populations and is modulated by the total initial cell density. A well-mixed model with competition mediated by resource sharing and cooperation from global degradation of toxins predicts the experimentally observed behavior. These observations suggest underlying population-level mechanisms that are largely independent of biofilm spatial structure.

## Introduction

Bacterial communities respond to the changing environments they inhabit. Take for example the dynamic behaviors and complex mechanisms by which bacteria respond to antibiotic stress, driven by processes operating across molecular, cellular, and population scales. A particularly relevant response is antibiotic resistance, which emerges not only from genetic determinants, but also from phenotypic traits and collective mechanisms ***Denk-Lobnig and Wood (2023***); ***Moran and Wood (2025***); ***Blair et al. (2015***); ***Corona and Martinez (2013***); ***Vega and Gore (2014***); ***Pearl Mizrahi et al. (2023***). Given the prevalence of antibiotic resistance and the variety of responsible mechanisms, understanding how they interact to produce community level responses is a necessity for a complete understanding of resistance in dynamic bacterial populations across their different growth modalities. We aim to study these community level responses in multi-strain populations of antibiotic resistant and sensitive bacteria in two different growth modalities–biofilms and their plankton subpopulation–while recording survival and dynamics in both.

A prevalent mechanism of antibiotic resistance is enzymatic drug degradation, by which enzymes produced by resistant bacteria irreversibly inactivate antibiotics ***Wright (2005***). This mechanism enables collective resistance and thus creates cooperative interactions that could allow sensitive cells to survive in reduced antibiotic concentrations. This phenomena has been observed in surface-attached biofilms as well as planktonic communities. In mixed planktonic cultures, resistant subpopulations rescue sensitive cells ***Yurtsev et al. (2013***); ***Sorg et al. (2016***); ***Pathak et al. (2023***); ***Gross et al. (2024***). In biofilms, additional intrinsic defense mechanisms (e.g., reduced antibiotic diffusion due to the extracellular matrix, and metabolic heterogeneity) further increase antibiotic resistance relative to planktonic populations ***Davey and O’toole (2000***); ***Donlan (2002***); ***Stewart and William Costerton (2001***); ***Flemming et al. (2016***); ***Hall and Mah (2017***); ***Liu et al. (2024***); ***Ruelens et al. (2025***). However, the intrinsic spatial organization of biofilms raises the question of whether the role of enzymatic-based cooperative resistance is amplified in these communities.

As a model organism, we employ *Enterococcus faecalis*, a non-motile Gram-positive bacterium that is opportunistically pathogenic. *E. faecalis* is a common cause of hospital acquired infections and frequently exhibits multidrug resistance ***Gilmore et al. (2014***). Furthermore, *E. faecalis* readily forms robust biofilms. Our mixed populations consist of a resistant strain of *E. faecalis* that produces β-lactamase enzymes, which degrade β-lactam antibiotics. The produced β-lactamase is not released from *E. faecalis* ***Murray (1992***), but resistant cells still act as antibiotic sinks, providing protection to surrounding bacteria rather than directly secreting a public good. This drug-dependent cooperation produces complex spatiotemporal patterns in agar-grown *E. feacalis* colonies composed of mixed sensitive and resistant bacteria ***Denk-Lobnig and Wood (2025***); ***Sharma and Wood (2021***). To more fully explore *E. faecalis*’s cooperative dynamics in the presence of antibiotic stress, we combine β-lactam antibiotic (ampicillin) with a *β*-lactamase inhibitor (sulbactam). In this work, we focus on understanding the effects of resistance and cross-protection in simultaneously growing, coupled biofilm and planktonic populations over a range of initial resistance levels. Modifying the initial densities, drug concentrations, and drug mixtures allows us to elucidate the effect of community structure on sensitive and resistant population dynamics. Unexpectedly, we find that antibiotic driven compositional shifts are the same in both biofilm and planktonic populations. Correlation analysis further revealed that the biofilms lacked clear spatial structuring, despite expecting spatial organization from local antibiotic sinks created by resistant cells.

## Results

### Resistant subpopulations protect biofilm and planktonic populations equally

To investigate the role of cooperative resistance in shaping bacterial response to antibiotic stress, we analyzed coupled biofilm and planktonic populations consisting of mixed ampicillin-sensitive and ampicillin-resistant *E. faecalis* bacteria. Both strains constitutively express distinct fluorescent markers—colored here as green for sensitive and magenta for resistant ***Hallinen et al. (2020a***). Resistance is conferred by a plasmid-encoded *β*-lactamase of clinical origin ***Hallinen et al. (2020b***). For each experiment, mixtures were prepared at specific initial percentages of resistant cells in an otherwise sensitive population (0%, 1%, 5%, 10%, 15%, 25%, and 50%) and two different cellular densities (optical density equal to 0.01 and 0.001). These populations were then grown in a glass bottom petri dish with liquid growth media supplemented with defined concentrations of antibiotics (0, 1, 2 or 3 µg/ml of ampicillin; 1 µg/ml of ampicillin with 1 or 2 µg/ml of sulbactam). After 24 hours of undisturbed incubation at 37°C, bacteria form two distinct populations, a surface-attached biofilm and a free-floating planktonic community. The final composition and characteristics of both populations were measured via confocal fluorescent microscopy Figure 1 (a). For details, refer to Materials and Methods.

**Figure 1.**
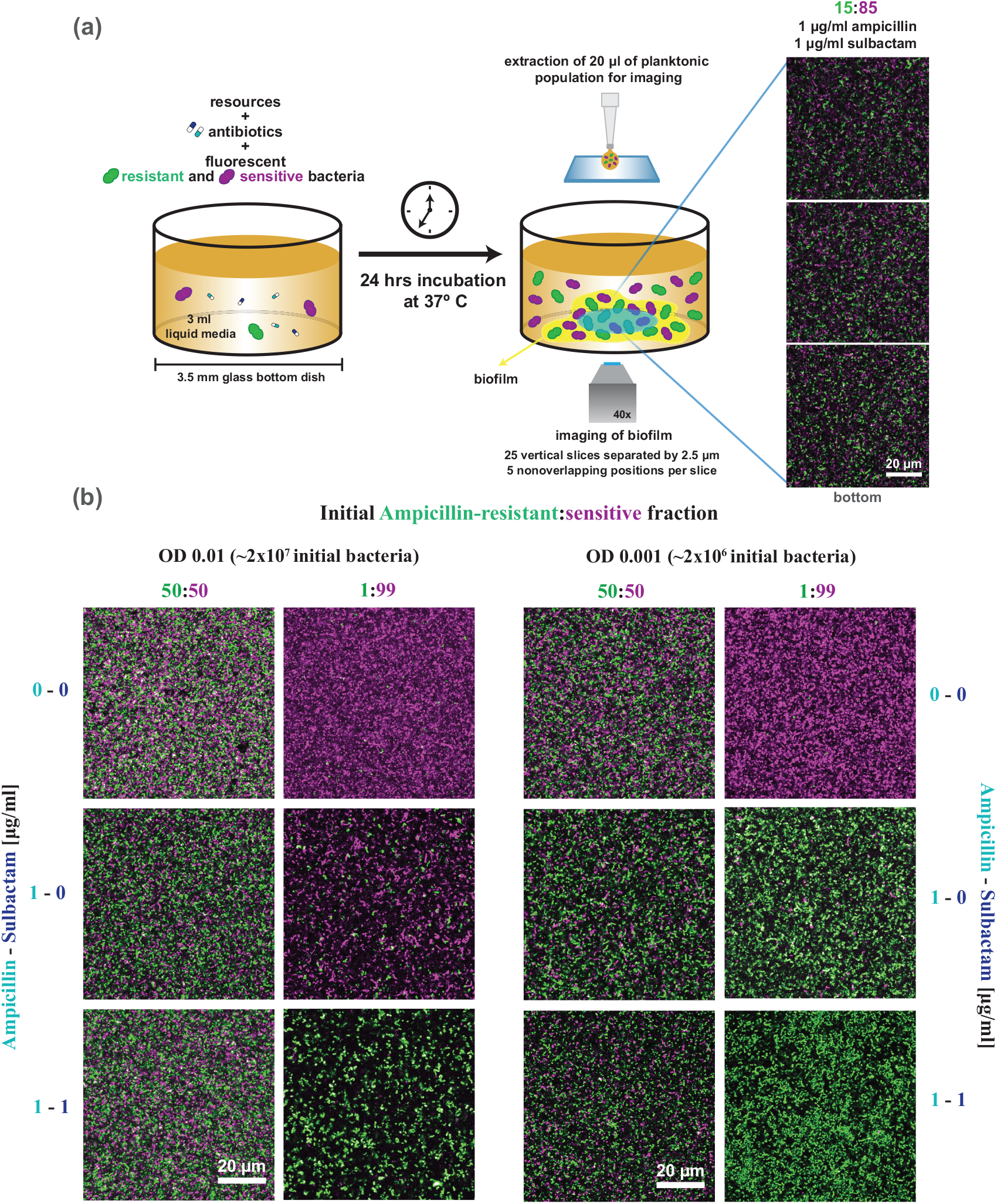
*E. faecalis* populations growing in the presence of ampicillin and sulbactam undergo significant compositional changes that depend on drug concentration, initial population density, and composition. (a) Diagram illustrating the experiment. A mixture of florescent ampicillin-sensitive and ampicillin-resistant *E. faecalis* bacteria, prepared at a defined density and ratio, is added to a 3.5 mm glass bottom dish containing 3 ml liquid growth media supplemented with drugs. After 24 hours of incubation at 37°C the biofilm is imaged directly in the glass bottom dish using confocal fluorescence microscopy to obtain a z-stack through the biofilm (25 cross-sectional slices of dimensions 80 µm x 80 µm with 2.5 µm of vertical separation). Z-stacks were obtained from five non-overlapping positions within the biofilm. The three slices depicted correspond to a biofilm with starting composition of 15% resistant and 85% sensitive bacteria treated with ampicillin at 1 µg/ml and sulbactam at 1 µg/ml. A sample of the planktonic population is extracted and mounted on a glass slide for imaging using confocal fluorescent microscopy. (b) Representative images (densest slice) of mixed ampicillin-resistant (green) and ampicillin-sensitive (magenta) biofilms after the incubation period. Initial bacterial density, initial resistant:sensitive ratio, as well as ampicillin and sulbactam concentrations were varied to examine effects on final population composition. The first row corresponds to untreated biofilms, the second to biofilms grown with ampicillin at 1 µg/ml, and the third to biofilms grown with both ampicillin and sulbactam at 1 µg/ml. All panels are shown at the same scale.

The presence of resistant strains in bacterial populations has been observed to protect non-*β*-lactamase producers from otherwise lethal ampicillin concentrations in diverse growth scenarios ***Nicoloff and Andersson (2016***); ***Frost et al. (2018***); ***Sharma and Wood (2021***); ***Denk-Lobnig and Wood (2025***); ***Yurtsev et al. (2013***); ***Amanatidou et al. (2019***); ***Sorg et al. (2016***); ***Ruelens et al. (2025***). This phenomena is evident in our biofilm microscopy data Figure 1(b), where, despite population-wide compositional shifts, sensitive bacteria (magenta) are always present in cultures with otherwise inhibitory concentrations of ampicillin. A sensitive subpopulation is still seen even with the addition of our *β*-lactamase inhibitor, sulbactam Figure 1(b, bottom row). In addition to the biofilms, we further examined the planktonic population in the supernatant and found sensitive cells were present there as well. We quantify the fraction of resistant bacteria in the final population as a function of its initial value for both biofilm and planktonic populations, revealing a non-monotonic dependency for all ampicillin concentrations tested, Figure 2. We refer to this as a population inversion effect, where, below a threshold value, the final fraction of resistant cells is greater than its initial value, leading to the increase of the resistant fraction in the population. This effect results from the interplay between two bacterial “social” interactions, competition for resources and cooperative protection against antibiotics. When the initial number of resistant bacteria is low, cross-protection against antibiotics is limited, sensitive bacteria are unable to survive, and resistant bacteria grow to a larger fraction of the final population. On the other hand, when the initial number of resistant bacteria is sufficiently large, sensitive cells are protected from the antibiotic and compete with the resistant cells for resources, limiting the resistant population. As a result, the population inversion effect is amplified at higher initial drug concentrations and attenuated at higher initial cell densities.

**Figure 2.**
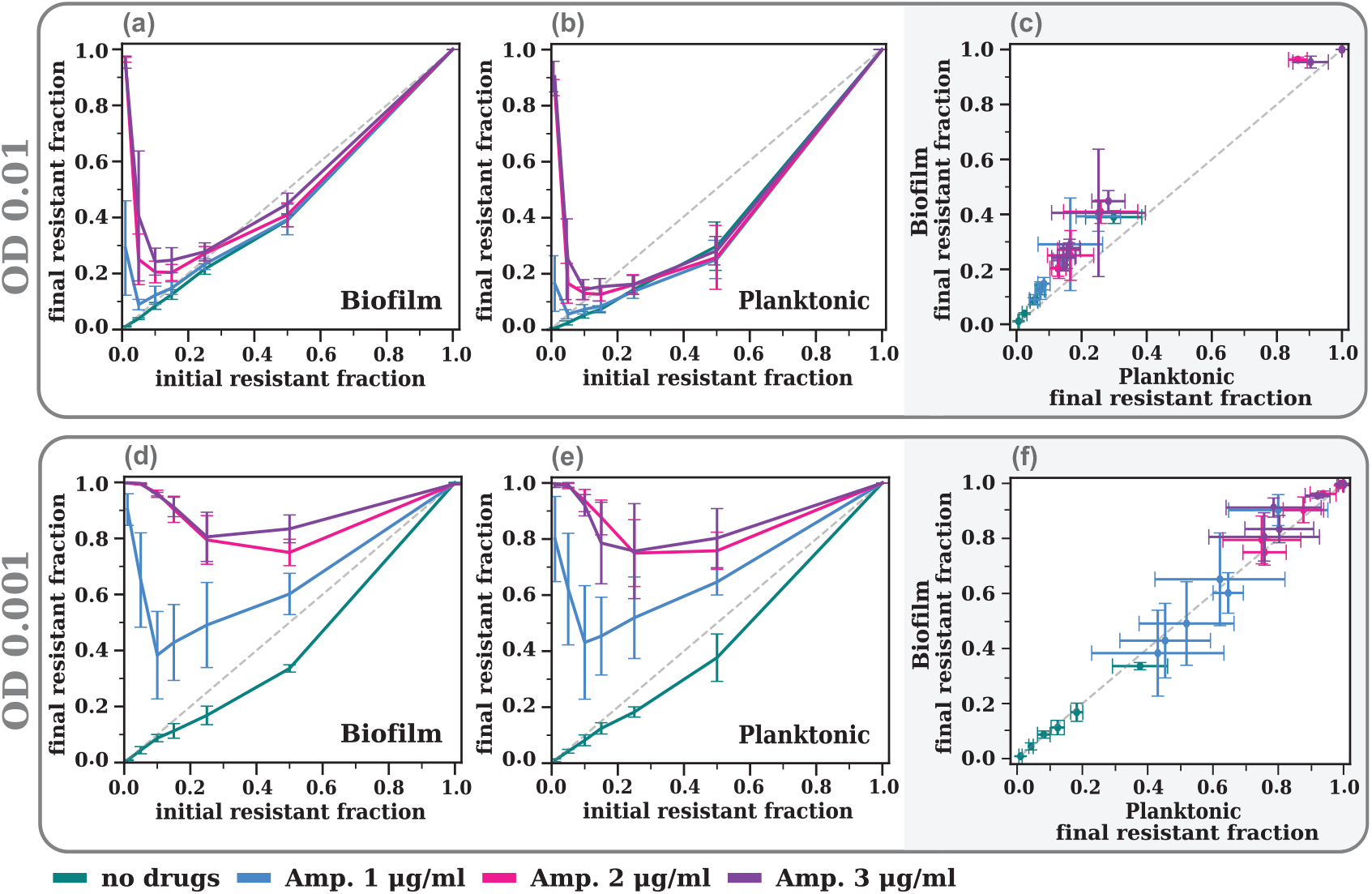
Biofilm and planktonic populations exhibit a non-monotonic dependence between final and initial resistant fractions after antibiotic treatment, with final composition of biofilm and planktonic populations being nearly identical. Rows indicate different initial starting OD’s. (a) & (d) Final resistant fraction versus initial resistant fraction in the biofilm for the no drugs case (green), and after treatment at Amp. 1 µg/ml (blue), Amp. 2 µg/ml (pink), and Amp. 3 µg/ml (purple). (b) & (e) Final resistant fraction versus initial resistant fraction in the planktonic. (c) & (f) Final resistant fractions in the biofilm versus final resistant fractions in the planktonic for the no drugs case and all Amp. concentrations. Error bars throughout all panels represent standard error across the five non-overlapping regions of the biofilm and the three experimental replicas.

### *β*-lactamase inhibition enhances resistance effects

To increase efficacy against strains exhibiting resistance mechanisms, *β*-lactam antibiotics are commonly administered in conjunction with *β*-lactamase inhibitors such as sulbactam. Sulbactam has little intrinsic antibacterial activity, but binds irreversibly to *β*-lactamases, thus protecting ampicillin from enzymatic degradation and restoring antibiotic efficacy ***Drawz and Bonomo (2010***); ***Bush (1988***); ***Carcione et al. (2021***); ***English et al. (1978***). Since resistant cells mediate protection of sensitive cells through *β*-lactamase activity, we expect the addition of sulbactam to alter the cooperative dynamics between the two strains.

As expected, sulbactam by itself does not alter the population composition relative to the drug-free case, Figure 3 panels (a) and (d). However, when combined with ampicillin, the final fraction of resistant cells is consistently higher than with ampicillin alone Figure 3, panels (b) and (e). This can be understood through the interplay of cooperative and competitive interactions, as the efficacy of the *β*-lactamase that the resistant cells produce is limited due to the inhibition by sulbactam, leading to decreased protection for the sensitive cells. The early loss of sensitive cells reduces competition for nutrients, allowing the fraction of resistant cells to rise within the population. As was observed in the ampicillin-only condition, biofilm and planktonic populations have indistinguishable compositions of sensitive and resistant cells Figure 3 panels (c) and (f).

**Figure 3.**
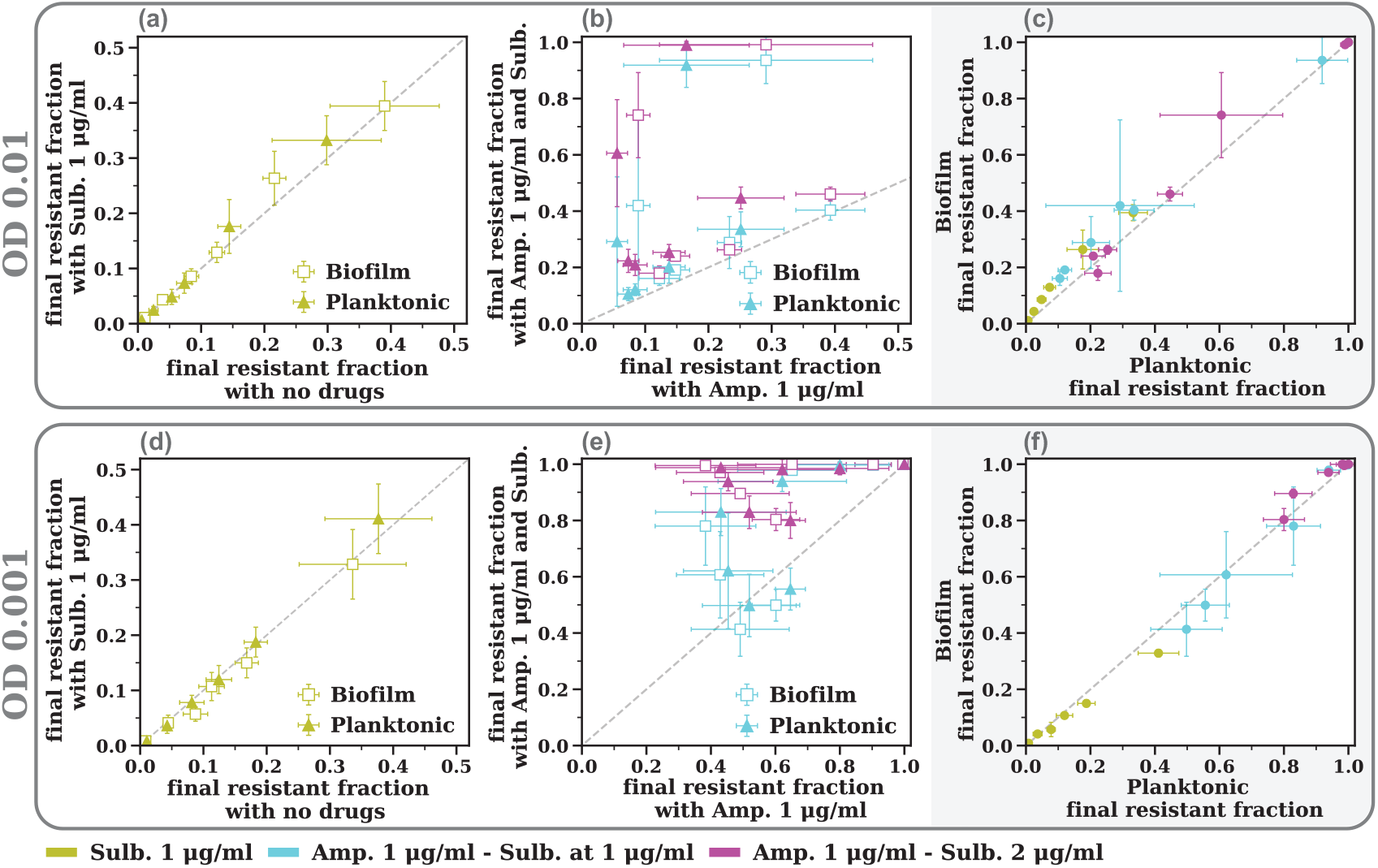
The combination of ampicillin and sulbactam increases the final resistant fraction in both biofilm and planktonic populations and leads to a nearly identical final composition. Rows indicate different initial starting OD’s. (a) & (d) Final resistant fraction for sulbactam experiments versus final resistant fraction for no drug experiments for both biofilm (square) and planktonic (triangle). (b) & (e) Final resistant fraction for ampicillin and sulbactam experiments versus final resistant fraction of just ampicillin experiments (blue: Amp µg/ml-Sulb µg/ml, purple: Amp µg/ml-Sulb. 2 µg/ml). (c) & (f) Final resistant fractions in the biofilm versus final resistant fractions in the planktonic for the Sulb. 1 µg/ml case and at the two cases with drug combinations. Error bars throughout all panels represent variation across the five non-overlapping regions of the biofilm and the three experimental replicas.

### Correlation analysis reveals biofilms are structurally amorphous

In addition to the characteristics of the global population of the planktonic and biofilm communities, we also analyzed the spatial structures of the biofilms. Previous work has noted biofilms often exhibit well-defined spatial structure that can be altered in response to antibiotic treatment ***Dale et al. (2017***). The local nature of this cooperative detoxification suggests that biofilms formed under antibiotic stress may exhibit distinct spatial structures, reflected in differing relative distances between the resistant *β*-lactamase producers and sensitive non-producers. We use a metric similar to a cumulative radial distribution function to study the distribution of resistant cells in relation to sensitive cells. The quantification of this metric, which we refer to simply as correlation, requires two datasets. One contains the extracted values of the centroid for each cell in the biofilm and the other is a randomized dataset that contains the same number of cells, but with centroid values assigned at random. The correlation is then equal to the ratio between the average number of resistant cells within a distance *r* of a sensitive cell in the original dataset and the same quantity calculated in the randomized dataset ***Stewart et al. (2013***); ***Martínez-García et al. (2018***); ***Schillinger et al. (2012***). When equal to one, this metric indicates that cells are positioned in the biofilm as if they were random. Correlation values below one indicate fewer resistant cells in the neighborhood of sensitive cells than expected in a random spatial arrangement, whereas values greater than one indicate more resistant cells than expected.

For distances larger than 1 µm, the correlation is equal to one, and drops below for distances smaller than 1 µm. This observation holds for biofilms formed with and without antibiotic stress for all values of drug concentrations and initial resistant fractions Figure 4. Given that the typical cell size is approximately 1 µm, values below one are expected, as only the sensitive cell of interest is present within this region. We find the biofilm structure to be largely random, showing no signature of spatial correlation. In conjunction with the observation that planktonic and biofilm communities undergo the same population shifts, these results suggest that the drug-dependent cooperative interaction does not lead to structure in our *E. faecalis* bacterial communities.

**Figure 4.**
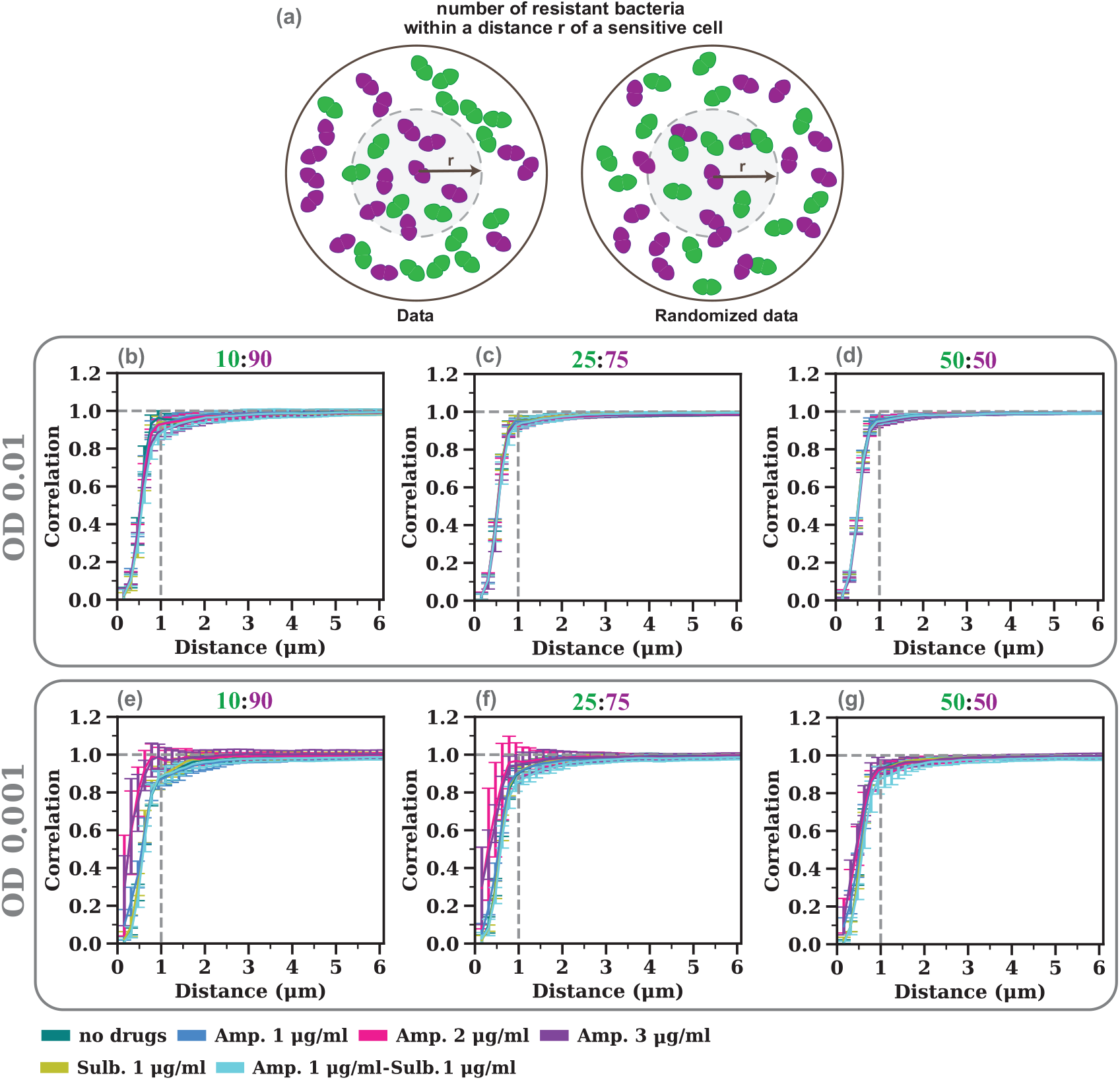
Correlation analysis reveals that the biofilm does not exhibit spatial structure. (a) Schematic of the correlation metric used to quantify structure: the ratio of the number of resistant bacteria within a distance r of a sensitive bacteria in the original data to that in the randomized data. Values equal to one indicate that the biofilm lacks spatial structure, as the relative positions are the same as in a completely random scenario. Values below one indicate a decrease in the number of resistant cells close to sensitive cells in relation to the random scenario. (b), (c) & (d) Correlation values for three different initial resistant fractions (10%, 25% & 50%) at an initial OD of 0.01. (e), (f) & (g) Correlation values for three different initial resistant fractions (10%, 25% & 50%) at an initial OD of 0.001. Dashed line at 1 µm indicates approximate cell length. Error bars represent variation across the five non-overlapping regions of the biofilm and the three experimental replicas.

### Well-mixed model captures experimental observations

Experimental data show that planktonic and biofilm populations have very similar final compositions, with no spatial structure in the biofilm. These observations suggest that a well-mixed model without spatial dynamics may be sufficient to reproduce the observed non-monotonic relationship between initial and final resistant fractions (Figures 2 and 3). To this end, we consider a model for the dynamics of the resistant (*R*_*t*_) and sensitive (*S*_*t*_) cell biomass composed of three core processes: resource limited cell birth, drug induced cell death, and drug degradation by resistant cells.

Growth is described using the empirical Monod model ***Monod (1949***); ***Koch (1998***); ***Shao et al. (2017***), with growth limiting glucose concentration *g*_*t*_. Furthermore, both strains undergo only ampicillin-dependent cell death, as sulbactam showed negligible toxicity at the concentrations tested. Thus, we take a death rate that depends on the ampicillin concentration *a*_*t*_ and not on the sulbactam concentration *s*_*t*_, which is assumed to have the form of a Hill function ***Regoes et al. (2004***). Together these observations lead to the collection of dynamical equations:

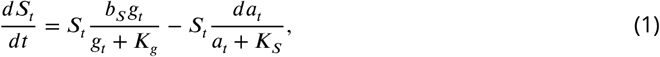

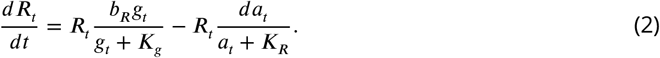

These equations incorporate the fitness cost of antibiotic resistance, as sensitive and resistant cells are assumed to have distinct maximum growth rates, *b*_*S*_ and *b*_*R*_, but with the same half-maximum glucose value *K*_*g*_. As sensitive cells are substantially more susceptible to ampicillin than resistant cells, the death rate of the sensitive and resistant strains reach half of the maximum rate *d* at different ampicillin concentrations, *K*_*S*_ and *K*_*R*_.

Throughout the experiment, the concentrations of glucose, ampicillin and sulbactam only decrease, as cells consume resources and intracellular *β*-lactamase degrades the antibiotics. With the biomass yield per unit of glucose consumed *Y*, we get the following equations:

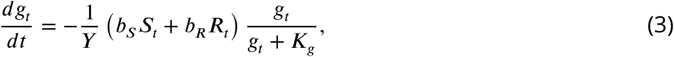

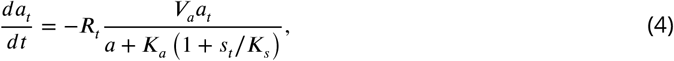

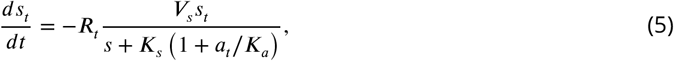

As both ampicillin and sulbactam bind to the same substrate, we take the binding rate of the drugs to *β*-lactamase to follow Michaelis-Menten kinetics with competitive inhibition ***Shapiro (2017***). The maximum degradation rates per cell are *V*_*a*_ and *V*_*s*_ with the presence of sulbactam increasing the effective Michaelis-Menten constant *K*_*a*_ for ampicillin by a factor of (1 + *s*_*t*_/ *K*_*s*_), and vice versa for sulbactam.

In this model, the dynamics proceed until the initial concentration of glucose (*g*_0_) is consumed and all the antibiotics (*a*_0_ and *s*_0_) are degraded. We assume that the resulting time-independent fixed point captures the final observed biofilm composition. This fixed point only depends on six dimensionless parameters that we choose based on the biophysical interpretations:

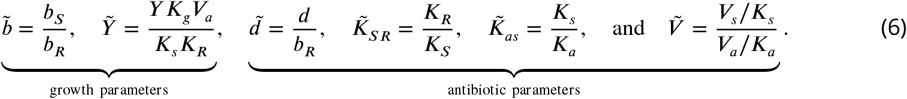

The first two determine the growth dynamics. They quantify the fitness difference between the strains, 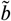, and a dimensionless yield coefficient, 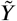 The remaining parameters determine the effects of the antibiotics through the relative rate of birth and death, 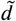, the difference in lethal antibiotic concentrations, 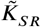, the relative binding strength of the antibiotics, 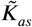, and the relative antibiotic degradation rates, 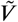. The solution to the model is then completely determined from the nondimensionalized initial biomass, 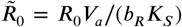and 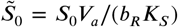, as well as the initial glucose, 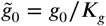, ampicillin, 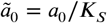, and sulbactam 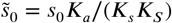, concentrations.

To validate the model, we proceed by fitting the six dimensionless parameters and the initial conditions to experimental data at the smaller initial OD of 0.001. This is done in sequence to check for internal consistency. First, in the absence of drug, the dynamics only depend on the growth parameters, 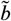 and 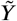, initial biomass 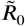 and 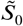, and initial glucose concentration, 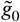. Thus, we can estimate these parameters using data only from the no-drug condition. Specifically, we use data on the number of sensitive cells to determine 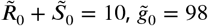, and 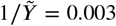. Refer to Materials and Methods for the data. The value of 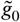 indicates that the system operates well above the saturating glucose concentration for growth. This is consistent with the nutrient-rich growth medium used. We then switch to using the resistant fraction data to achieve an improved fit for the ratio of birth rates, 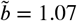, supporting the hypothesis that resistance incurs a fitness cost.

With these parameters in hand, we can proceed to determine the parameters related to druginduced cell death. Using the resistant fraction data for the lowest ampicillin concentration (Amp. 1 µg/ml) we estimate the values of 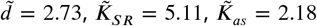, and 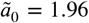, while holding all the previously determined growth parameters fixed. Finally, data for the lowest sulbactam concen-tration (Amp. 1 µg/ml–Sulb. 1 µg/ml) is used to fit the remaining parameters, 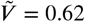 and 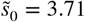. The value of *K*_*SR*_ is consistent with the expectation that resistant bacteria reach their maximum death rate at higher ampicillin concentrations than sensitive cells. The small value of 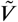 reflects the higher efficiency of ampicillin degradation compared to sulbactam.

We next challenge the model by evaluating its predictions on the subset of data not used for fitting. Specifically, we compare measurements at higher ampicillin concentrations (Amp. 2 µg/ml and Amp. 3 µg/ml) with model predictions generated using twofold, 2*ã*_0_, and threefold, 3*ã*_0_, increases of the fitted initial ampicillin concentration. We also compare data at the highest initial OD with model predictions generated using a ten-fold increase of the fitted initial cell biomass, 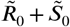. For both values of initial cell densities the model reproduces the observed nonmonotonic behavior and is quantitatively accurate except for the highest ampicillin concentration, Figure 5, panels (b) and (d). In the presence of sulbactam (Sulb. 1 µg/ml, Amp. 1 µg/ml - Sulb. 1 µg/ml and Amp. 1 µg/ml - Sulb. 2 µg/ml), model predictions were generated using the fitted baseline ampicillin and sulbactam concentrations with multiplicative scaling where appropriate. Although the model predicts an increase in the final fraction of resistant cells relative to the cases with ampicillin alone, it does not quantitatively capture the observed values. This could reflect additional interactions between the drugs that are not included in the model.

**Figure 5.**
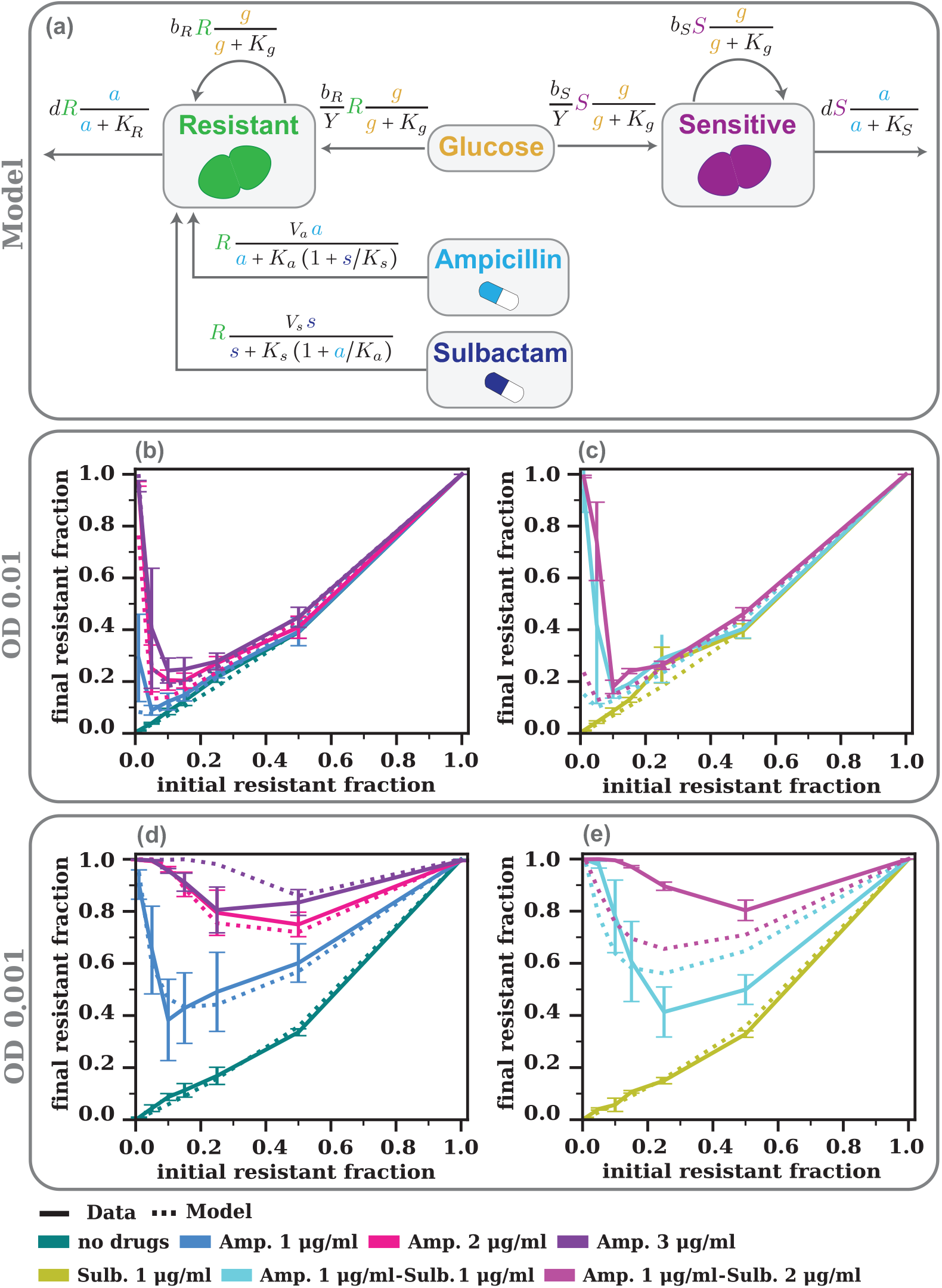
(a) Representation of the model. Sensitive and resistant bacteria grow as they consume glucose, and die with rates that depend on the ampicillin concentration. Resistant bacteria degrade both ampicillin and sulbactam. (b) & (c) Comparison between experimental data (solid lines) and model prediction (dashed lines) for the final resistant fraction across all drug combinations at an initial OD of 0.01. (d) & (e) Comparison between experimental data and model predictions, including fitting, at an initial OD of 0.001. In panel (d), the no drugs and Amp. 1 µg/ml curves are fitted to extract model parameters. All other curves shown in the figure are model predictions.

## Discussion

Bacterial communities are shaped by both their environment and their responses to that environment. Studying a mixed bacterial population composed of *E. faecalis* antibiotic sensitive and resistant cells, we were able to examine these interactions and dynamic processes in both the surfaceattached biofilm and the planktonic community. Given the specific form of antibiotic resistance– the production of *β*-lactamase enzymes–we expected to observe cooperative detoxification, and our studies were able to fully capture this effect in both the attached biofilm and well-mixed plank-tonic populations. We find that both planktonic and biofilm populations undergo a population inversion effect under antibiotic treatment, characterized by final resistant fractions greater than their starting value. Somewhat surprisingly, this effect is quantitatively indistinguishable between biofilm and planktonic populations despite their different environmental contexts. The impact of the resistant cells–and by extension the *β*-lactamase enzymes those cells produce–is modulated by the initial cell density, where the initial number of resistant cells affects the drug degradation dynamics and therefore the amount of cross-protection provided to sensitive cells. Studies using planktonic cultures of *Escherichia coli* found similar protection for sensitive cells when growing with a resistant sub-population, suggesting cooperative protection is a broadly applicable phenomenon ***Yurtsev et al. (2013***); ***Ruelens et al. (2025***). However, previous work found that population shifts caused by antibiotic stress were quantitatively distinct in planktonic and biofilm cultures of *E. coli* grown separately ***Amanatidou et al. (2019***). However, unlike *E. faecalis, E. coli β*-lactamase is found extracellularly, naturally altering the mechanistic details of the cooperative interaction ***Francisco et al. (1992***).

In addition to the overall population composition, the confocal microscopy data allowed us to specifically explore the spatial organization and individual single-cell positioning within biofilms. Instead of finding signal of spatial correlations within the biofilm (e.g., sensitive cells localized more frequently near resistant cells), we instead found that the spatial organization was largely random. This disparity could reflect differences in the enzyme location mechanism or indicate that the simultaneous formation of the two populations during antibiotic stress produces distinct effects than when the two populations are studied separately. The lack of spatial organization that we observe experimentally suggests that the population inversion effect can be explained by employing a well-mixed model with no spatial extension. Fitting a model of cooperative interactions between resistant and sensitive strains mediated by ampicillin degradation and resource consumption with data from only the lowest ampicillin concentration surprisingly allowed us to reproduce the quantitative and qualitative aspects of the remaining experimental data.

## Acknowledgments

In memory of Professor Kevin Wood, who inspired a generation of scientists. GFM would like to thank all members of the Wood lab for helpful discussions and assistance with experiments. GFM and JMH acknowledge support from the National Science Foundation under Grant No. 2142466. LZ acknowledges support from NIH grant RM1 DE034220.

## Materials and Methods

### Bacterial strains

Experiments were performed with *E. faecalis* antibiotic resistant and sensitive OG1RF strains, fluorescently labeled with a plasmid that encoded a spectinomycin resistance gene. Sensitive cells carried PBSU101-DasherGFP, while resistant cells carried PBSU101-blaZ-RudolphRFP. Both plasmids were constitutively expressed. The blaZ gene was derived from the CH19 *E. faecalis* clinical isolate ***Rice et al. (1991***). The design and construction of the reporter plasmids were previously described in ***Hallinen et al. (2020a***,b). Brain Heart Infusion broth (Remel) was used throughout the experiment as the growth media. Selection of the plasmids was enforced by supplementing all media with spectinomycin sulfate (Thermo Fisher Scientific, at 120 µg/ml).

### Biofilm growth experiment

Overnight cultures were prepared by inoculating bacteria directly from frozen stocks into media and grown at 32 °C on an orbital shaker at 150 RPM. After 17 hours, cultures were diluted at a 1:4 ratio to enable for accurate optical density (OD_600_) measurements. Both cultures were brought to an OD_600_ of 0.51. Mixtures with a target initial percentage of resistant bacteria equal to *p* were prepared by mixing 2 ×*p* µl of the resistant culture and 2 ×(100 ™*p*) µl of the sensitive culture in 10 ml of media. For experiments starting at an OD of 0.001, an additional 1:10 dilution was performed. A total of 2.7 ml of each mixture was transferred to 35 mm glass bottom microscope dishes (Cellvis, D35-20-1-N) and incubated at 37°C. Samples were kept stationary during the incubation period. After 30 minutes, 0.3 ml of media supplemented with antibiotics was added to each dish. The antibiotics used include ampicillin sodium salt (Sigma-Aldrich) and sulbactam (Cayman Chemical). To achieve a target antibiotic concentration of *c* in 3 ml, the media supplemented with antibiotics was prepared at a concentration equal to 10 ×*c*. Samples were then incubated at 37°C for 24 hours.

### Data collection and image analysis

After 24 hours, the glass bottom dishes containing the biofilms were removed to room temperature and imaged separately on a Zeiss LSM 700 inverted confocal microscope with a 40x oil objective and a pinhole of 1.2 µm. Biofilms were imaged directly on the dish without prior washing. Five nonoverlaping z-stacks of dimensions 80 µm x 80 µm x 60 µm were imaged for each biofilm. Each z-stack was composed of 25 vertical slices separated by 2.5 µm and selected to allow for imaging of the biofilm from its bottom layer to its upper surface at every point. The composition of the planktonic population was measured by imaging three independent 20 µl samples on a glass slide. After image processing using Fiji ***Schindelin et al. (2012***), data analysis was performed in two steps, with pixel classification executed with scientific software ilastik ***Berg et al. (2019***) followed by bacterial segmentation using a custom MATLAB script.

### Model fitting

The resistant fraction data alone are insufficient to uniquely determine all growth parameters. To address this limitation, we additionally consider measurements of the total number of sensitive cells in the biofilm as a function of the initial resistant fraction, Figure 6.

**Figure 6.**
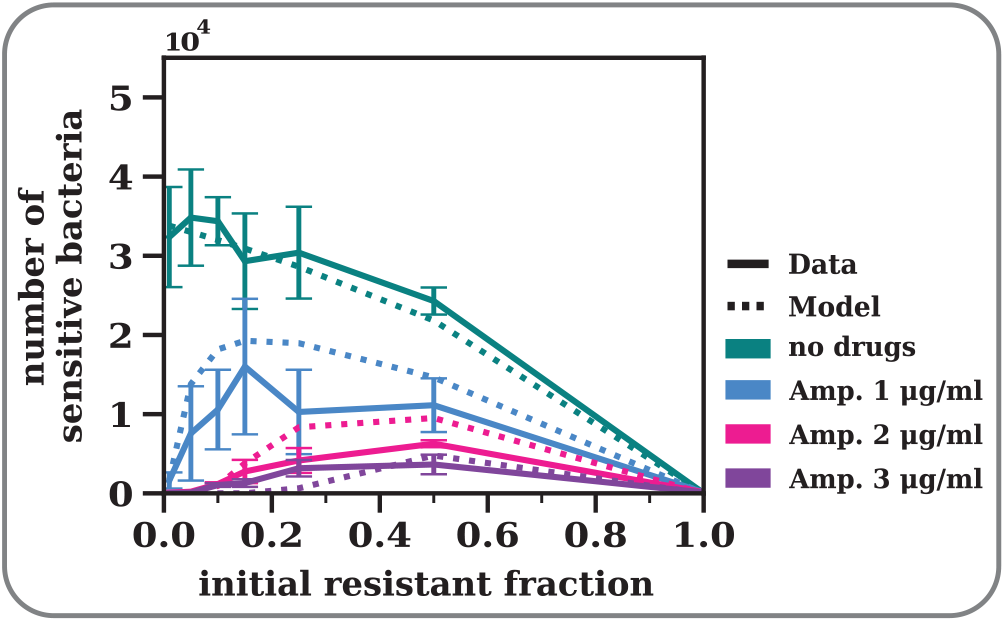
Experimental data (solid lines) and model predictions (dashed lines) for the total number of sensitive bacteria within a biofilm z-stack at initial OD 0.001. The data in the absence of drugs (green) is used to fit the growth parameters 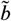and 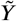, as well as the initial total biomass and glucose concentrations. Predictions for the lowest ampicillin concentration (blue) are generated using the parameter values obtained from the fitting procedure. Predictions at Amp. 2 µg/mL (pink) and Amp. 3 µg/mL (purple) are obtained by increasing the fitted initial ampicillin concentration twofold and threefold, respectively.

